# Biochemical characterization of the sole *Leptospira interrogans* diadenylate cyclase

**DOI:** 10.1101/2025.04.15.648992

**Authors:** Edward J.A. Schuler, Dhara T. Patel, Richard T. Marconi

## Abstract

Leptospirosis, a widespread zoonosis impacting humans, companion animals, livestock, and wildlife, is caused by a diverse group of *Leptospira* species. Leptospires can infect virtually any vertebrate and can survive in and adapt to radically different environments. Adaptive responses are dependent on a sophisticated network of sensing and signaling systems. Bioinformatic analyses revealed that *Leptospira* species encode a single CdaA-type diadenylate cyclase (DAC). DACs catalyze the synthesis of cyclic di-adenosine monophosphate (c-di-AMP) from two molecules of ATP. The regulatory roles of c-di-AMP in *Leptospira* have not yet been explored. Herein, we present a biochemical analysis of the *L. interrogans* str. Fiocruz L1-130 CdaA-type diadenylate cyclase, LIC10844. Triton X-114 extraction, phase partitioning, and subsequent immunoblot analyses revealed that LIC10844 is an inner membrane-associated protein. Recombinant CdaA was produced and tested for DAC activity and cofactor specificity. Cobalt and manganese were competent divalent cations, whereas zinc, magnesium, copper, nickel, and calcium were not. Notably, the optimal concentrations of cobalt and manganese for maximal enzymatic activity differed significantly. Size exclusion chromatography and subsequent DAC assays of the eluate revealed the enzymatically active form of CdaA to be a homodimer. Amino acid residues involved in DAC activity were identified through site-directed mutagenesis and DAC assays. This report is the first to characterize a DAC in *Leptospira*. The data suggest that *Leptospira* utilizes c-di-AMP as a second messenger molecule and that its enzymatic activity is responsive to environmental divalent cation concentrations.

## Introduction

Leptospirosis, the most common zoonosis worldwide, poses a significant health threat to humans, companion animals, livestock, and wildlife (1). The causative agents are a group of antigenically diverse *Leptospira* species. There are estimated to be over one million human cases of leptospirosis each year, resulting in nearly 60,000 deaths (2). However, due to misdiagnosis and the lack of reporting infrastructure, the actual number of cases is estimated to be much greater (3). The disease may be self-limiting or progress to a life-threatening multi-system disorder affecting the kidneys, liver, and lungs (4). Diverse animals have been demonstrated to be competent reservoir hosts. Leptospires establish infection of the renal tubules and are shed in urine. Infection can occur through direct contact with the urine or indirect contact with contaminated water or soil.

Pathogenic *Leptospira* species encode over 50 genes annotated as components of two-component regulatory systems and transcriptional regulators. The expansive repertoire of potential regulatory proteins allows for rapid sensing and adaptation to environmental stimuli (5, 6). Cyclic nucleotides have been demonstrated to be important secondary messenger molecules that directly or indirectly regulate diverse cellular processes. Among bacteria and archaea, c-di-AMP is the most widespread second messenger molecule. Here, we investigate the functional activity of the diadenylate cyclase (DAC), CdaA, the sole DAC predicted to be encoded by all *Leptospira* species. DACs catalyze the formation of c-di-AMP from two molecules of ATP (7, 8). The effector mechanisms of c-di-AMP in *Leptospira* species remain undefined.

Crystal structures of orthologous CdaA proteins identified an ATP binding site and two active site motifs: “DGA” and “GXRHRXA” (9). Herein, we demonstrate that CdaA is conserved amongst diverse pathogenic *Leptospira* species and expressed by all isolates tested. The enzymatic activity of CdaA is dependent on the presence and concentration of specific divalent cations. Single-amino acid substitutions within the putative active and ATP-coordination sites attenuated enzymatic activity. The active form of CdaA was determined to be a homodimer. The results of this study suggest that *Leptospira* utilizes c-di-AMP as a second messenger molecule and that its activity is responsive to environmental variables, including divalent cation availability and concentration.

## Materials and Methods

### Phylogenetic analyses

*Leptospira* CdaA amino acid sequences retrieved from NCBI (10) were aligned using MEGA11 (11), and a neighbor-joining tree (100 bootstraps; Poisson amino acid substitution model; pairwise deletion of gaps) was constructed. CdaA orthologues from *B. subtilis* and *L. monocytogenes* served as outliers. Percent amino acid identity and similarity values were calculated using the “Ident and Sim” program (12).

### Bacterial cultivation

*Leptospira* isolates (Table 1) were cultivated (30 °C) in EMJH medium supplemented with Probumin vaccine grade bovine serum albumin (BSA; EMD Millipore) and 100 µg mL^-1^ of 5-fluorouracil, as previously described (13). Growth was monitored by dark-field microscopy. Cells were recovered by centrifugation and washed twice with phosphate-buffered saline (PBS; 137 mM NaCl, 2.7 mM KCl, 2 mM KH_2_PO_4_, 1 mM Na_2_HPO_4_; pH 7.4).

**Table 1.** Bacterial isolates utilized in this study.

| **Isolate** | **Source** |
| --- | --- |
| *Leptospira interrogans* sv. Copenhageni str. Fiocruz L1-130 | Human, Brazil |
| *Leptospira interrogans* sv. Canicola str. Kito | Canine, Brazil |
| *Leptospira interrogans* sv. Pomona str. LC82-25 | Human, USA |
| *Leptospira interrogans* sv. Manilae str. L495 | Human, Philippines |
| *Leptospira noguchii* sv. Autumnalis str. Bonito | Human, Brazil |
| *Leptospira noguchii* sv. Australis str. Hook | Canine, Brazil |
| *Leptospira noguchii* sv. Bataviae str. Cascata | Human, Brazil |
| *Leptospira kirschneri* sv. Grippotyphosa str. RM52 | Swine, USA |
| *Leptospira borgpetersenii* sg. Ballum str. M2 | Mouse, Puerto Rico |
| *Leptospira kmetyi* sv. Malaysia str. Bejo-Iso9T | Soil, Malaysia |
| *Leptospira wolffii* sv. Korat str. Korat-H2T | Human, Thailand |

#### Generation and purification of recombinant proteins

Genes encoding CdaA or CdaA site-directed mutants were synthesized fee-for-service (GenScript) and supplied in pET-45b(+). The sequence encoding the putative N-terminal 87 amino acid transmembrane domain was omitted from all constructs based on DeepTMHMM predictions (15). Plasmids were propagated in *E. coli* DH5α cells (New England Biolabs), recovered with a QIAprep Spin Miniprep kit (QIAGEN), and sequence verified on a fee-for-service basis. (GeneWiz). Using standard methods, E. coli BL21 (DE3) cells (New England Biolabs) were transformed with the plasmids. Protein expression and purification were performed as previously described (14). In brief, protein expression was induced with isopropyl β-D-1-thiogalactopyranoside (IPTG; 1 mM). Cells were recovered by centrifugation and resuspended in binding buffer (500 mM NaCl, 20 mM Na_2_HPO_4_, 20 mM imidazole; pH 7.4) supplemented with 1% protease inhibitor cocktail (PIC; Sigma-Aldrich) and 2.5 U mL^-1^ of Pierce^™^ Universal Nuclease (Thermo Scientific). Cells were lysed using an EmulsiFlex-C3 high-pressure homogenizer (3 passes, 1000-1500 psi, 4 °C). The soluble fraction was recovered by centrifugation, and the recombinant proteins were purified by nickel affinity chromatography on an ÄKTA Pure 25 M FPLC (Cytiva) using 1 mL HisTrap FF columns (Cytiva) (16). Protein-containing eluates were pooled and dialyzed into PBS. Protein concentrations were determined by bicinchoninic acid assay (Pierce™), and purity was assessed by SDS-PAGE.

### SDS-PAGE and immunoblot analyses

Washed *Leptospira* cells were resuspended in 1:1 PBS:2X SDS-PAGE buffer (20% glycerol, 4% SDS, 2% β-mercaptoethanol, 0.1% bromophenol blue (w/v), 250 mM Tris (pH 6.8)) to a final concentration of 3.33 OD_600_ mL^-1^. Cells were lysed by sonication and heating (10 min; 99 °C). Recombinant proteins were prepared for SDS-PAGE by dilution in 2X SDS-PAGE buffer to a final concentration of 100 ng μL^-1^. Samples were heated (10 min; 99 °C), and 5 μL separated in Criterion™ TGX™ AnykDa gels (Bio-Rad) with Tris/Glycine/SDS Buffer (Bio-Rad), followed by staining with Coomassie brilliant blue R-250 (Thermo Scientific). Precision Plus Protein™ Dual Color Standards (Bio-Rad) served as the molecular weight (MW) markers. Stained proteins were imaged on a ChemiDoc™ Touch imaging system (Bio-Rad) using auto-optimal settings. Proteins were transferred to polyvinylidene difluoride (PVDF) membranes using the Trans-Blot® Turbo™ system (Bio-Rad; high MW preset). Membranes were blocked (15 min) in EveryBlot blocking buffer (Bio-Rad). Primary antibodies (1:1000 in blocking buffer) and goat anti-rat IgG HRP secondary antibody (Novus Biologicals; 1:40000 in blocking buffer) were incubated with blots for 1 hr each. After incubation with antibodies, the membranes were washed with PBST (PBS with 0.2% Tween-20 three times (5 min). Clarity Western ECL Substrate (Bio-Rad) was added (5 min), and images were captured as above.

### Generation of hyperimmune serum

Anti-CdaA_88-273_ antiserum was generated in Sprague-Dawley rats (Charles River) as previously described (17). In brief, prime intraperitoneal vaccination consisted of 50 µg CdaA_88-273_ (1:1 Freund’s complete adjuvant; Sigma-Aldrich), followed by two booster doses of 25 µg protein (2 weeks apart; 1:1 in Freund’s incomplete adjuvant; Sigma-Aldrich). Upon termination (CO_2_ inhalation followed by cervical dislocation), blood was collected by cardiac puncture, and serum harvested using VACUETTE Z serum sep clot activator tubes (Greiner Bio-One). All animal studies were conducted in accordance with the Guide for the Care and Use of Laboratory Animals (Institute of Animal Research, National Research Council) with protocols approved by the Virginia Commonwealth University IACUC.

### Triton™ X-114 extraction and phase partitioning

Triton™ X-114 extraction and phase partitioning were performed as previously described (18, 19), with some modifications. Approximately 10^10^ mid-log phase *L. interrogans* strain Fiocruz L1-130 cells were recovered by centrifugation and washed twice with wash buffer (5 mM MgCl_2_ in PBS pH 7.4). Cells were resuspended at 3.0 OD_600_ mL^-1^ in wash buffer, centrifuged, and finally resuspended in 1 mL of extraction buffer (1% Triton™ X-114, 1% PIC, 150 mM NaCl, 10 mM Tris-HCl (pH 7.5), 1 mM EDTA), and incubated on ice for 2 hr. The soluble and insoluble (protoplasmic cylinder; PC) fractions were separated by centrifugation. The supernatant was collected, and the PC was washed twice with 1 mL of extraction buffer and then stored dry at −20 °C. The supernatant was treated with 20 mM CaCl_2_, incubated (37 °C, 15 min), centrifuged to separate the aqueous phase (AQ) from the detergent-soluble phase (DET) and each washed twice with either 500 µL extraction buffer (AQ) or wash buffer (DET). Proteins in the AQ and DET phases were precipitated with 10 volumes of ice-cold acetone (rocking on ice; 30 min) and centrifuged. The acetone was evaporated, the phases were resuspended in 100 µL PBS, and the samples were analyzed by SDS-PAGE.

### Diadenylate cyclase (DAC) assays

DAC assays were performed as described (20), with some modifications. In brief, 5 μM recombinant protein and 150 μM ATP (New England Biolabs) were incubated in DAC buffer (50 mM Tris HCl, 50 mM NaCl, 0.5 mM EDTA with 10 mM MgCl_2_, MnCl_2_, CoCl_2_, ZnSO_4_, CuCl_2_, NiCl_2_, or CaCl_2_; pH 7.5) at 30 °C or 37 °C for 0.5, 1, 2, 4, 16, or 24 hr. After incubation, samples were heated at 95 °C for 5 min, centrifuged (15,200 x g; 2 min), filtered (0.22 μm PVDF membrane; EMD Millipore), and stored in amber fixed insert vials (Agilent) at 4 °C.

### Reverse phase high-performance liquid chromatography (RP-HPLC)

The RP-HPLC protocol utilized herein was synthesized from two previous publications (20, 21). DAC reactions and standards (150 μM ATP and c-di-AMP; Millipore Sigma) were separated in a Supelcosil LC-18-T 3 μm column (Millipore Sigma) on a 1260 Infinity II HPLC system (Agilent). Samples (20 μL) were injected into equilibrated columns. A linear gradient of Buffer A (100 mM KH_2_PO_4_, 4 mM tetrabutylammonium hydrogen sulfate; pH 5.9) and Buffer B (100% methanol) was generated, starting with 100% Buffer A, increasing Buffer B to 50% over 20 min at a flow rate of 0.7 mL min^-1^. Nucleotide retention times (Rt; min) were measured at 254 nm. The conversion efficiency for each sample was calculated based on area under the curve analyses compared to the c-di-AMP standard.

### Fluorescence-based detection of c-di-AMP

Fluorescence-based coralyne assays were used in parallel to RP-HPLC to measure the concentration of c-di-AMP in DAC reactions, as previously described (22). In brief, DAC assays were run as above. Master mixes (100 μL) were prepared as follows: 65% DAC reaction, 25% quenching buffer (1 M KBr, 150 mM NaCl, 25 mM Tris; pH 7.2), and 10% coralyne solution (150 mM NaCl, 25 mM Tris, 100 μM coralyne chloride (Sigma-Aldrich); pH 7.2). Samples were pipetted (30 μL) into a black, 384-well plate (Corning) in triplicate, mixed on an orbital shaker, then read on a BioTek Synergy H1 microplate reader at λ_ex_ = 420 nm (27 nm bandwidth) and λ_em_ = 540 (25 nm bandwidth). Concentrations of c-di-AMP in each DAC reaction were determined relative to the fluorescence of the c-di-AMP standard.

### Structural predictions of CdaA

The structure of LIC10844 was predicted with AlphaFold (23) using the ChimeraX application (24). The published crystal structure of CdaA from *Listeria monocytogenes* in complex with ATP and divalent cation (PDB code 4RV7) (9) was used to overlay the AlphaFold predicted structure of LIC10844 to calculate root mean square deviation (RMSD) in the PyMOL Molecular Graphics System, Version 4.6.0, Schrödinger, LLC. Specific residues forming polar contacts with ATP and/or the divalent cation were also predicted in PyMOL.

### Size exclusion chromatography (SEC)

SEC was performed as previously described (25), using a TSKgel G4000SW_XL_ column (Tosoh Bioscience) attached to a 1260 Infinity II HPLC system. PBS served as the mobile phase. A standard curve was generated using 15-600 kDa standard protein mix (Millipore Sigma). The standards were prepared per manufacturer instructions, filtered through a 0.22 µm PVDF membrane, injected into the column (20 μL), and eluted with PBS (flow rate of 1.0 mL min^-1^). Retention times (Rt; min) were plotted versus the Log(MW) to calculate a linear regression formula. To assess oligomeric state of CdaA proteins, 50 µg (in PBS) was filtered as above, injected into the column (25 μL), and eluted with PBS (flow rate of 1.0 mL min^-1^). Protein under the peaks (254 nm) was collected and concentrated for subsequent DAC assays.

### Statistical analyses

Ordinary one-way ANOVA followed by Dunnett’s multiple comparisons test, and two-way ANOVA fit with a full model were performed in GraphPad Prism 10.3.1 for Windows, GraphPad Software, Boston, Massachusetts USA, www.graphpad.com.

## Results

### CdaA is a conserved inner membrane protein

A CdaA neighbor-joining phylogenetic tree delineated two distinct clades consistent with the separation of pathogenic (P1/P2) from saprophytic (S1) *Leptospira* species (Figure 1). Pathogenic species group into P1 and P2 subclades (Figure 1), consistent with orthologous gene analyses (26). Percent amino acid identity values ranged from 88.7% to 99.0% across P1 strains and 72.0% to 74.6% among P2 strains. Relative to *L. interrogans*, saprophytic species encode more divergent CdaA homologs (53.6% to 54.7%).

**Figure 1.**
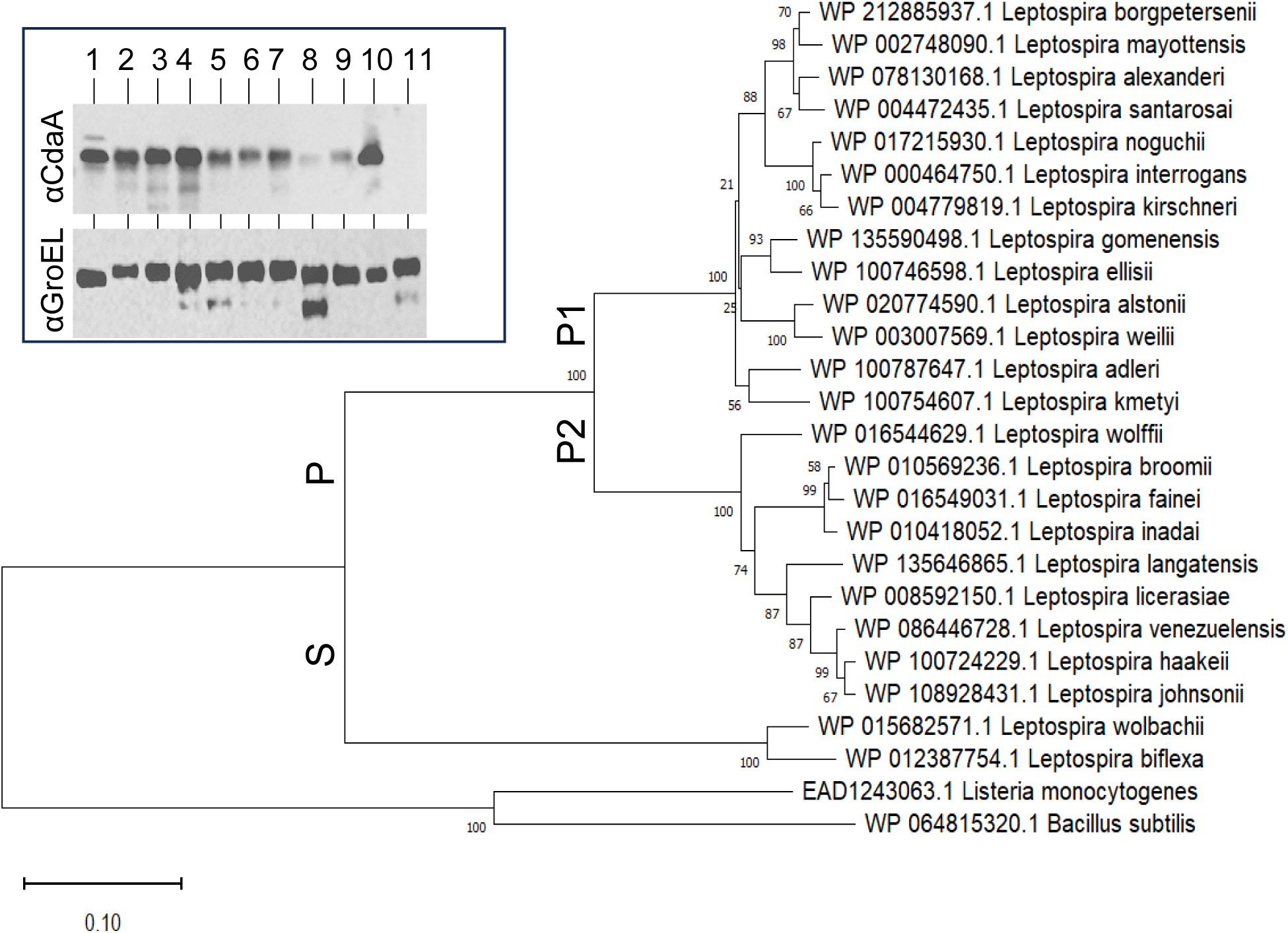
A neighbor-joining tree was constructed as detailed in the text. A distance scale is shown at the bottom left, and bootstrap frequencies are indicated at each branch. Pathogenic (P) and saprophytic (S) clades and the pathogenic (P1) and intermediate (P2) subclades are labeled at each branch point. The insert shows an immunoblot in which pathogenic isolates were screened with anti-CdaA_88-273_ and anti-GroEL (control) antisera, as described in the text. The isolates screened were: 1) *L. interrogans* str. Fiocruz L1-130, 2) *L. interrogans* str. Kito, 3) *L. interrogans* str. LC82-25, 4) *L. interrogans* str. L495, 5) *L. noguchii* str. Bonito, 6) *L. noguchii* str. Hook, 7) *L. noguchii* str. Cascata, 8) *L. kirschneri* str. RM52, 9) *L. borgpetersenii* str. M2, 10) *L. kmetyi* str. Bejo-Iso9T, and 11) *L. wolffii* str. Korat-H2T. Note that the immunoblots were cropped to remove empty space.

The production of CdaA during *in vitro* cultivation was assessed by immunoblot analysis of ten P1 strains and one P2 strain of *Leptospira* (Table 1). Cell lysates were screened with anti-CdaA_88-273_ antiserum and anti-GroEL antiserum as a loading control. As expected, GroEL (∼60 kDa) was detected in all strains (Figure 1 insert). CdaA (∼31 kDa) was detected in all strains except the P2 clade strain, *L. wolffii* str. Korat-H2T. The percent amino acid identity value for the CdaA proteins of *L. interrogans* and *L. wolffii* is 73%. The lack of detection by the antiserum likely reflects antigenic divergence as opposed to a lack of expression during cultivation. The subcellular location of CdaA was determined through immunoblot analysis of fractions obtained from Triton™ X-114 extraction and phase partitioning. Immunoblots were screened with anti-CdaA_88-273_ antiserum. Screening with anti-LipL32 and anti-GroEL antiserum served outer membrane and cytoplasmic controls, respectively (Figure 2). CdaA localized to the protoplasmic cylinder (Figure 2) exclusively. This observation and the presence of three predicted transmembrane domains in the N-terminus of the protein support the conclusion that CdaA is an inner membrane protein.

**Figure 2.**
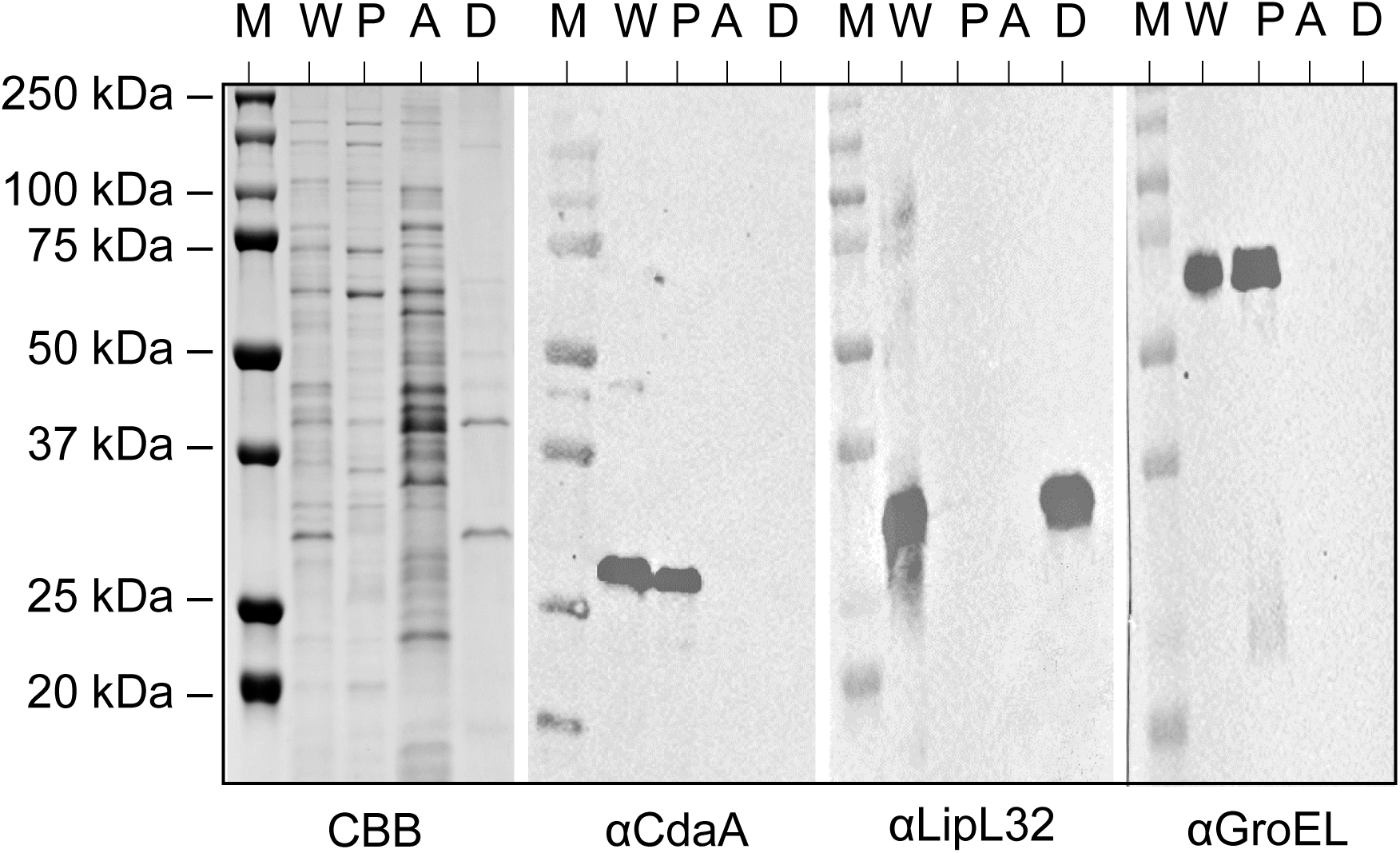
Localization of CdaA by Triton X-114 extraction and phase partitioning. *L. interrogans* str. Fiocruz L1-130 cells were extracted with Triton X-114 and phase partitioned as described in the methods. A Coomassie brilliant blue (CBB) stain of the whole cell lysate (W), periplasmic cylinder (P), aqueous (A), and detergent-soluble (D) phases is shown. Immunoblots were screened with anti-CdaA_88-273_, anti-LipL32, and anti-GroEL antiserum, as indicated below each image. The migration position of a molecular weight marker (M) is shown on the left.

### CdaA is a functional diadenylate cyclase

To assess the DAC activity and divalent cation cofactor requirements of CdaA, a fragment spanning residues 88-273 was produced. Attempts to express full-length CdaA were unsuccessful due to three predicted transmembrane domains within the first 87 amino acids of the protein. CdaA_88-273_ was expressed in *E. coli* BL21 (DE3) cells at high levels and was successfully purified to homogeneity. DAC activity was assessed over time in the presence of 10 mM MnCl_2_ at either 30 °C or 37 °C to identify optimal incubation conditions for subsequent assays. There was no significant difference between CdaA conversion efficiency at 30 °C versus 37 °C; however, maximum conversion of ATP to c-di-AMP (91% ± 4%) was not reached until 16 hr (Figure 3 A). To examine specific divalent cofactor requirements of CdaA, DAC assays were conducted in the presence of Mn^2+^, Mg^2+^, Co^2+^, Zn^2+^, Cu^2+^, Ni^2+^, or Ca^2+^. Only Mn^2+^ and Co^2+^ served as competent cofactors (Figure 3 B). Notably, significantly less cobalt (2.5 mM) was required than manganese (5 mM) for maximal conversion efficiency (Figure 3 C). Furthermore, CdaA was functional with as little as 1 mM cobalt, versus a minimum of 2.5 mM manganese required (Figure 4 B), highlighting the potential utility of the rarer cobalt cofactor in nutrient-poor environments where manganese may be unavailable.

**Figure 3.**
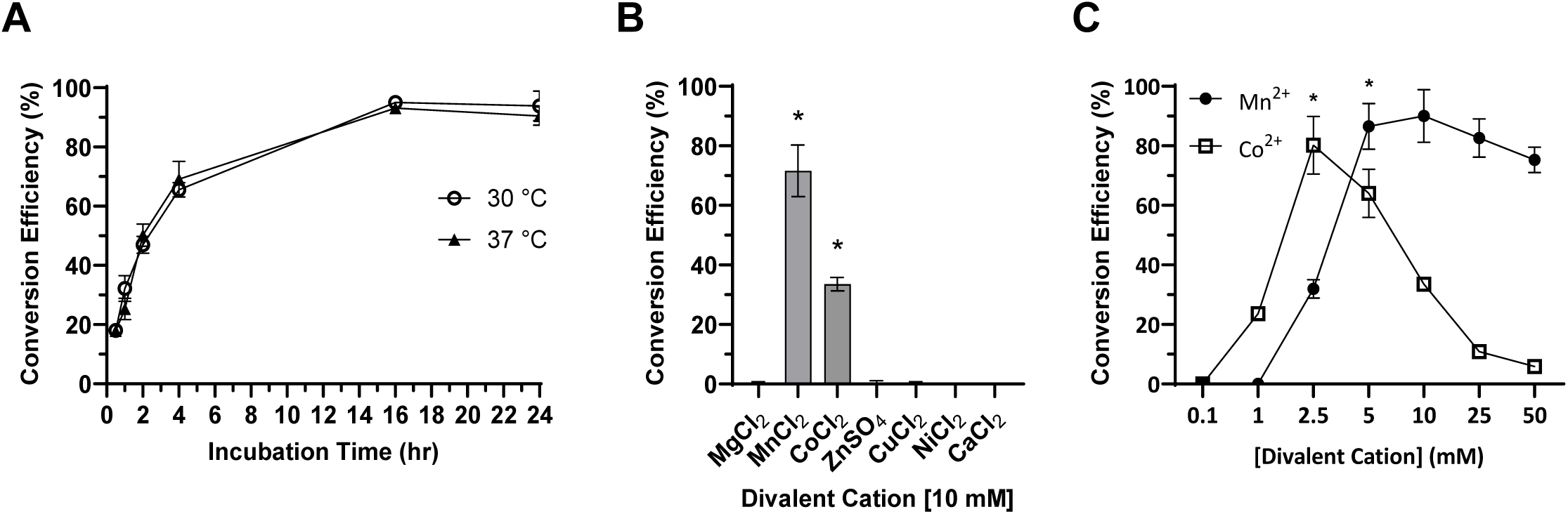
Optimization of diadenylate cyclase assay conditions. In panel A, conversion of ATP to c-di-AMP was assessed after 0.5, 1, 2, 4, 16, and 24 hr incubation with CdaA_88-273_ at 30 °C and 37 °C in 10 mM Mn2+ buffer. In panel B, the activity of CdaA_88-273_ in the presence of various divalent metal cations (10 mM) was assessed after 16 hr at 30 °C with ATP. Significance (P < .05) values are shown in comparison to buffer with ATP alone. Panel C shows the effect of divalent cation concentration on the enzymatic activity of CdaA_88-273_ after a 16 hr, 30 °C incubation with ATP. Asterisks represent the minimum required concentration of divalent cation to achieve maximal conversion efficiency (P < .05). All conversion efficiencies were calculated using coralyne fluorescence assays based on a c-di-AMP standard curve, as described in the text. One of three representative experimental replicates is shown, each with three technical replicates. Results were confirmed by RPC-HPLC area under the curve analyses, as described in the methods.

**Figure 4.**
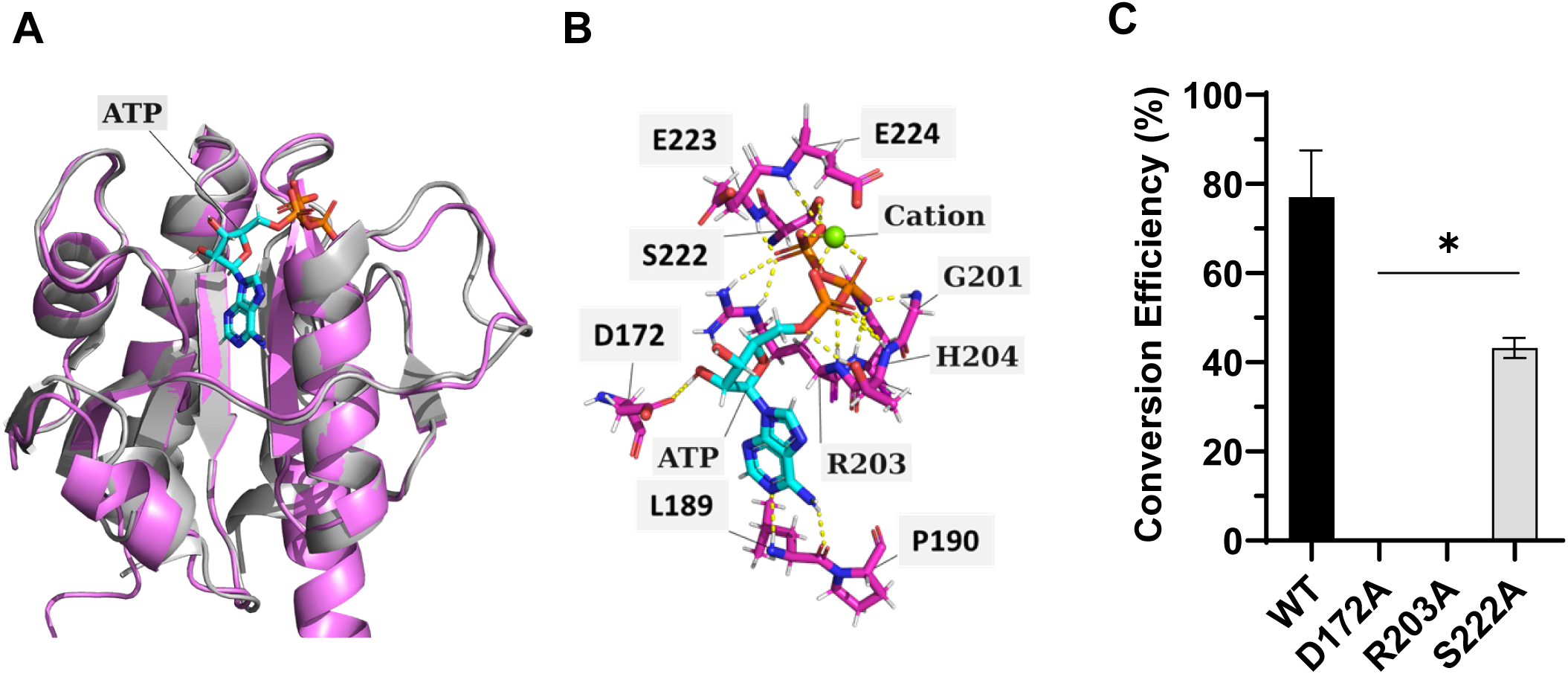
Identification of CdA residues required for DAC activity. The predicted AlphaFold structure of LIC10844 (pink) was overlayed on the crystal structure of the *Listeria monocytogenes* CdaA/ATP complex (gray; Panel A). CdaA residues predicted to form polar contacts with ATP or the divalent cation are shown in Panel B. The DAC activity of CdaA proteins harboring site-directed mutations was measured using a coralyne fluorescence-based assay as detailed in the text (Panel C). The proteins were incubated with ATP and 10 mM Mn^2+^ (30 °C; 16 hr), and conversion efficiencies were calculated as described in the text. Significance (* P < .05), relative to CdaA_88-273_ was determined by one-way ANOVA.

### Identification of CdaA residues required for DAC activity

Although no crystal structure is available for any leptospiral CdaA, the crystal structure of an orthologous CdaA from *L. monocytogenes* in a complex with ATP has been determined (9). The AlphaFold structural prediction of LIC10844 agreed closely with the crystal structure of *Listeria* CdaA+ATP (RMSD = 0.529) (Figure 4 A), despite only 35.0% amino acid identity and 52.1% amino acid similarity between the two proteins. Therefore, polar contacts with ATP or the divalent cation in the *Listeria* CdaA+ATP complex were identified and extrapolated to the corresponding residues in LIC10844 (Figure 4 B). Three residues were selected for initial mutational analyses: Asp-172 from the “<u>D</u>GA” active site, Arg-203 from the “GX<u>R</u>HRXA” active site, and Ser-222, a residue predicted to coordinate the ATP and the divalent cation. Three mutated recombinant LIC10844 proteins were generated, each with a single-alanine substitution in one of the aforementioned residues. DAC activity of each was measured after 0.5, 1, 2, 4, and 16 hr at 30 °C using RP-HPLC and coralyne fluorescence assays. Substitutions of Asp-172 or Arg-203 with alanine rendered the protein non-functional (Figure 4 C). Interestingly, the S222A mutant was still functional but demonstrated significantly reduced conversion efficiency (47.5% ± 8.1%) at 16 hr (Figure 4 C).

### The active form of CdaA is a homodimer

Recombinant CdaA_88-273_ and the three single-point mutants (D172A, R203A, and S222A) were separated by size exclusion chromatography and eluant under each peak collected for further analyses. The retention times of CdaA88-273 and each of the three single-point mutant proteins were identical. Two peaks with retention times (Rts) of ∼ 5.5 and 9.1 min were observed (Figure 5 A). A MW standard curve was generated as described in the methods to generate a linear regression formula: Log(MW) = −0.3546×(Rt(min)) + 4.843 (R^2^ = 0.9860). Based on the standard curve, the protein under peak 1 has an MW = 781.1 kDa, representing a recombinant protein aggregate. The calculated MW for the protein under peak 2 is 41.3 kDa in size, consistent with a homodimer (predicted MW of the His-tagged CdaA_88-273_ monomer = 22.1 kDa).

**Figure 5.**
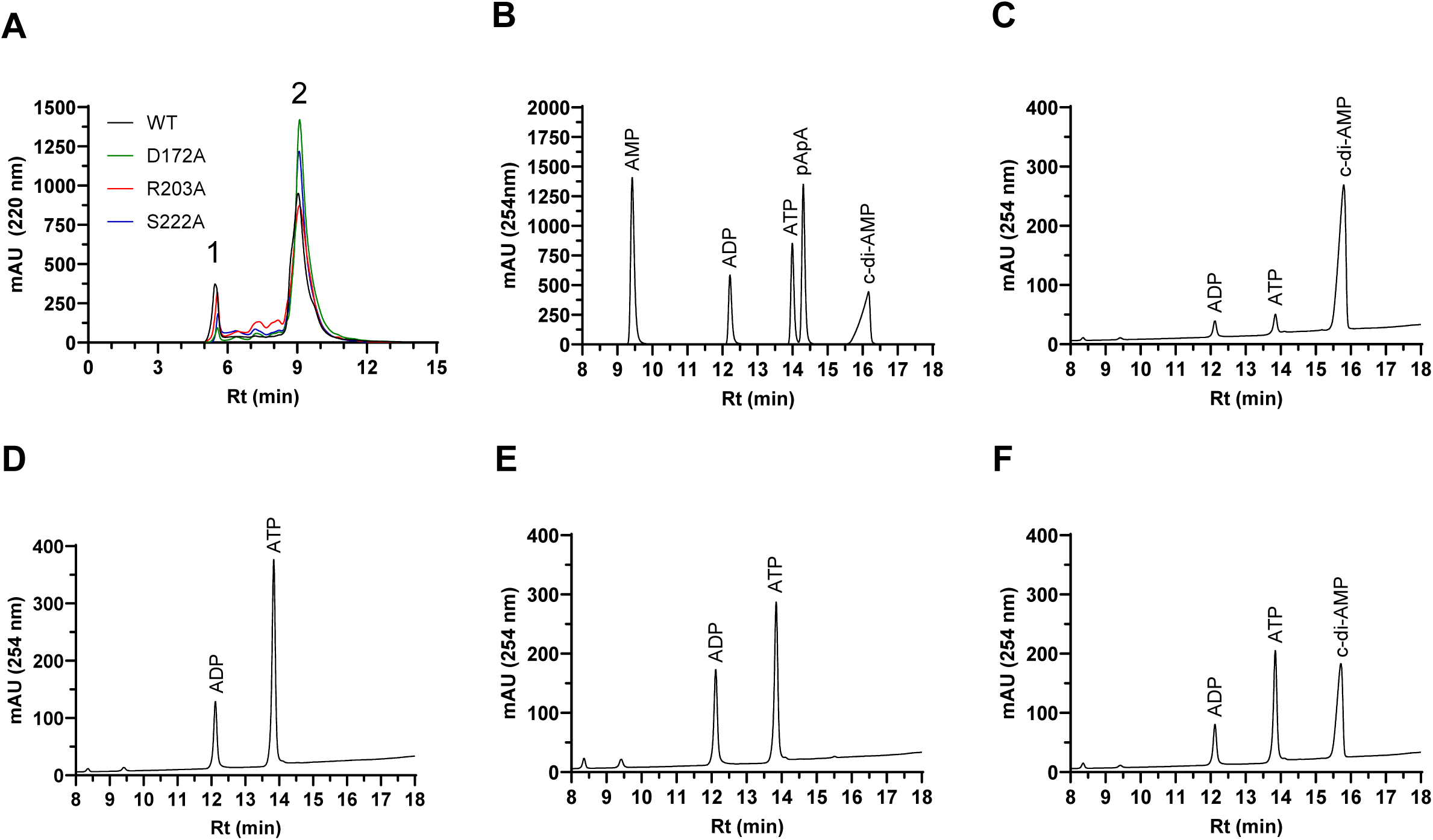
CdaA oligomerization and enzymatic activity. CdaA_88-273_ and the site-directed mutans were assessed using size exclusion chromatography (Panel A). Fractions under the peaks were collected. The protein under peak 2 was concentrated by acetone precipitation. Panel B shows nucleotide standards analyzed by RP-HPLC. Panels C through F show RPC-HPLC chromatograms of dimers (SEC peak 2) of CdaA_88-273_, CdaA_88-273_ D172A, CdaA_88-273_ R203A, and CdaA_88-273_ S222A after incubation with ATP (16 hr; 30 °C). The identity of each peak is indicated based on the nucleotide standards. Retention time (Rt; min) is presented on the x-axis. Representative data from one of three independent experiments are shown.

To determine if the homodimer is the enzymatically active form, the protein under each peak 2 was collected, concentrated, and tested for DAC activity. Dimeric CdaA_88-273_ from SEC peak 2 (Figure 5 A) was enzymatically active, converting ∼ 83.5% of the input ATP to c-di-AMP after 16 hr (Figure 5 C). As seen previously, dimeric CdaA_88-273_ D172A and R203A were unable to convert ATP to c-di-AMP (Figure 5 D-E), and CdaA_88-273_ S222A demonstrated a reduced efficiency of 56.6% (Figure 5 F). These data demonstrate the formation of an enzymatically-active CdaA_88-273_ homodimer *in vitro*.

## Discussion

The regulatory role of the second messenger molecule, c-di-AMP, has not been investigated in the pathogenesis and biology of *Leptospira* species. This study identified a single, CdaA-type diadenylate cyclase encoded by pathogenic and saprophytic species of *Leptospira*. CdaA specifically requires Mn^2+^ or Co^2+^ for DAC activity but is incapable of utilizing Mg^2+^ and is active as a homodimer—similar to the CdaA protein of *L. monocytogenes* (8, 9). Conversely, DisA-type DACs have been shown to be active in the presence of Mg^2+^ (27, 28, 29) as homo-octamers (27). It has been proposed that the capability of pathogens to utilize cobalt ions is advantageous for survival in environments with limited nutrients and competition for divalent cations (9, 30).

The vertebrate host is one such example of a nutrient-competitive environment. As the host attempts to sequester divalent cations, pathogens that can utilize uncommon ions, such as cobalt, may have increased fitness (30). *Leptospira* species have been shown to invade and persist in nutrient-replete macrophages (31, 32). It has been suggested that the ability of CdaA to utilize Mn^2+^ or Co^2+^, but not Mg^2+^ like DisA (33), may be due to differences in the amino acid composition of the active site (9). There is a conserved histidine immediately before the “DGA” motif in LIC10844, also found in CdaA of *L. monocytogenes* (8), but not in DisA proteins, that has low affinity for Mg^2+^, but can still bind Mn^2+^ and Co^2+^ (34). However, *Staphylococcus aureus* CdaA, which harbors the conserved histidine, can utilize Mg^2+^. It is important to note that there are significant conformational differences in metal binding sites between of *S. aureus* and *L. monocytogenes* CdaA that may account for their differing ability to utilize Mg^2+^ (8, 35). It remains to be determined how divalent cation preferences regulate CdaA activity *in vivo* and contribute to the ability of *Leptospira* to persist in the environment and host.

While CdaA is known to be essential for virulence in the relapsing fever spirochete, *Borrelia turicatae* (36), such mutational studies have yet to be conducted in *Leptospira*. It is likely that pathogenic leptospires similarly depend on c-di-AMP as a second messenger molecule. *Leptospira* encode *cdaA* in a 19-gene operon that includes *cdaR* but not *glmM* (37). CdaR and GlmM have been identified across bacteria as CdaA-binding proteins that regulate c-di-AMP production (35, 38, 39). The lack of GlmM in the leptospiral CdaA operon suggests unique regulatory mechanisms. Further studies are necessary to elucidate these mechanisms in *Leptospira*.

This study is the first to investigate a leptospiral diadenylate cyclase. Herein, we determined that single-alanine substitutions of Asp-172 or Arg-203 rendered CdaA_88-273_ non-functional and that mutation of Ser-222 resulted in ∼40% reduced conversion efficiency compared to wild-type despite the ability of all three mutant proteins to dimerize. It was not unexpected that mutations of the conserved Asp-172 and Arg-203 residues would eliminate DAC activity, as they are both found within the putative “D^172^GA” and “GXR^203^HRXA” active sites. Rosenberg et al. solved the first crystal structure of a CdaA orthologue in complex with ATP in *L. monocytogenes* and found Ser-222 to interact with the β- and γ-phosphates of ATP (9). Our data shows that the loss of this ATP-coordination residue attenuates but does not abolish the conversion of ATP to c-di-AMP. Further mutational and structural analyses are required to elucidate the role of additional functional residues in dimerization and enzymatic activity.

In summary, pathogenic and saprophytic isolates of *Leptospira* encode a single, functional, CdaA-type diadenylate cyclase. The biologically active form of CdaA is a homodimer complexed with either manganese or cobalt. Our understanding of the role of c-di-AMP in leptospiral pathogenesis is limited and warrants further investigation.

## Acknowledgments

Dr. David Haake (University of California, Los Angeles) and Dr. Albert Ko (Yale University, New Haven) kindly provided the Leptospira isolates utilized in this study. We thank members of the Marconi lab for reviewing this manuscript. This work was supported in part by funds from the School of Medicine and the Department of Microbiology and Immunology at Virginia Commonwealth University.

## References

1. Adler B, de la Pena Moctezuma A. 2010. Leptospira and leptospirosis. Vet Microbiol 140:287–96.

2. Hagedoorn NN, Maze MJ, Carugati M, Cash-Goldwasser S, Allan KJ, Chen K, Cossic B, Demeter E, Gallagher S, German R, Galloway RL, Habuš J, Rubach MP, Shiokawa K, Sulikhan N, Crump JA. 2024. Global distribution of Leptospira serovar isolations and detections from animal host species: A systematic review and online database. Trop Med Int Health 29:161–172.

3. Costa F, Hagan JE, Calcagno J, Kane M, Torgerson P, Martinez-Silveira MS, Stein C, Abela-Ridder B, Ko AI. 2015. Global Morbidity and Mortality of Leptospirosis: A Systematic Review. PLoS Negl Trop Dis 9:e0003898.

4. Bharti AR, Nally JE, Ricaldi JN, Matthias MA, Diaz MM, Lovett MA, Levett PN, Gilman RH, Willig MR, Gotuzzo E, Vinetz JM. 2003. Leptospirosis: a zoonotic disease of global importance. The Lancet Infectious Diseases 3:757–771.

5. Nascimento AL, Ko AI, Martins EA, Monteiro-Vitorello CB, Ho PL, Haake DA, Verjovski-Almeida S, Hartskeerl RA, Marques MV, Oliveira MC, Menck CF, Leite LC, Carrer H, Coutinho LL, Degrave WM, Dellagostin OA, El-Dorry H, Ferro ES, Ferro MI, Furlan LR, Gamberini M, Giglioti EA, Góes-Neto A, Goldman GH, Goldman MH, Harakava R, Jerônimo SM, Junqueira-de-Azevedo IL, Kimura ET, Kuramae EE, Lemos EG, Lemos MV, Marino CL, Nunes LR, de Oliveira RC, Pereira GG, Reis MS, Schriefer A, Siqueira WJ, Sommer P, Tsai SM, Simpson AJ, Ferro JA, Camargo LE, Kitajima JP, Setubal JC, Van Sluys MA. 2004. Comparative genomics of two Leptospira interrogans serovars reveals novel insights into physiology and pathogenesis. J Bacteriol 186:2164–72.

6. Missiakas D, Raina S. 1998. The extracytoplasmic function sigma factors: role and regulation. Molecular Microbiology 28:1059–1066.

7. Fahmi T, Port GC, Cho KH. 2017. c-di-AMP: An Essential Molecule in the Signaling Pathways that Regulate the Viability and Virulence of Gram-Positive Bacteria. Genes (Basel) 8.

8. Heidemann JL, Neumann P, Dickmanns A, Ficner R. 2019. Crystal structures of the c-di-AMP-synthesizing enzyme CdaA. J Biol Chem 294:10463–10470.

9. Rosenberg J, Dickmanns A, Neumann P, Gunka K, Arens J, Kaever V, Stülke J, Ficner R, Commichau FM. 2015. Structural and biochemical analysis of the essential diadenylate cyclase CdaA from Listeria monocytogenes. J Biol Chem 290:6596–606.

10. O’Leary NA, Wright MW, Brister JR, Ciufo S, Haddad D, McVeigh R, Rajput B, Robbertse B, Smith-White B, Ako-Adjei D, Astashyn A, Badretdin A, Bao Y, Blinkova O, Brover V, Chetvernin V, Choi J, Cox E, Ermolaeva O, Farrell CM, Goldfarb T, Gupta T, Haft D, Hatcher E, Hlavina W, Joardar VS, Kodali VK, Li W, Maglott D, Masterson P, McGarvey KM, Murphy MR, O’Neill K, Pujar S, Rangwala SH, Rausch D, Riddick LD, Schoch C, Shkeda A, Storz SS, Sun H, Thibaud-Nissen F, Tolstoy I, Tully RE, Vatsan AR, Wallin C, Webb D, Wu W, Landrum MJ, Kimchi A, et al. 2016. Reference sequence (RefSeq) database at NCBI: current status, taxonomic expansion, and functional annotation. Nucleic Acids Res 44:D733–45.

11. Kumar S, Stecher G, Li M, Knyaz C, Tamura K. 2018. MEGA X: Molecular Evolutionary Genetics Analysis across Computing Platforms. Mol Biol Evol 35:1547–1549.

12. Stothard P. 2000. The sequence manipulation suite: JavaScript programs for analyzing and formatting protein and DNA sequences. Biotechniques 28:1102, 1104.

13. Matsunaga J, Sanchez Y, Xu X, Haake DA. 2005. Osmolarity, a key environmental signal controlling expression of leptospiral proteins LigA and LigB and the extracellular release of LigA. Infect Immun 73:70–8.

14. Izac JR, Oliver LD, Jr., Earnhart CG, Marconi RT. 2017. Identification of a defined linear epitope in the OspA protein of the Lyme disease spirochetes that elicits bactericidal antibody responses: Implications for vaccine development. Vaccine 35:3178–3185.

15. Hallgren J, Tsirigos KD, Pedersen MD, Almagro Armenteros JJ, Marcatili P, Nielsen H, Krogh A, Winther O. 2022. DeepTMHMM predicts alpha and beta transmembrane proteins using deep neural networks. bioRxiv doi:10.1101/2022.04.08.487609:2022.04.08.487609.

16. Mallory KL, Miller DP, Oliver LD, Jr., Freedman JC, Kostick-Dunn JL, Carlyon JA, Marion JD, Bell JK, Marconi RT. 2016. Cyclic-di-GMP binding induces structural rearrangements in the PlzA and PlzC proteins of the Lyme disease and relapsing fever spirochetes: a possible switch mechanism for c-di-GMP-mediated effector functions. Pathog Dis 74.

17. Izac JR, O’Bier NS, Oliver LD, Jr., Camire AC, Earnhart CG, LeBlanc Rhodes DV, Young BF, Parnham SR, Davies C, Marconi RT. 2020. Development and optimization of OspC chimeritope vaccinogens for Lyme disease. Vaccine 38:1915–1924.

18. Schuler E, Marconi RT. 2021. The Leptospiral General Secretory Protein D (GspD), a secretin, elicits complement-independent bactericidal antibody against diverse Leptospira species and serovars. Vaccine X 7:100089.

19. Haake DA, Chao G, Zuerner RL, Barnett JK, Barnett D, Mazel M, Matsunaga J, Levett PN, Bolin CA. 2000. The leptospiral major outer membrane protein LipL32 is a lipoprotein expressed during mammalian infection. Infect Immun 68:2276–85.

20. Ryjenkov DA, Tarutina M, Moskvin OV, Gomelsky M. 2005. Cyclic diguanylate is a ubiquitous signaling molecule in bacteria: insights into biochemistry of the GGDEF protein domain. J Bacteriol 187:1792–8.

21. Li H, Li T, Zou W, Ni M, Hu Q, Qiu X, Yao Z, Fan J, Li L, Huang Q, Zhou R. 2022. IPA-3: An Inhibitor of Diadenylate Cyclase of Streptococcus suis with Potent Antimicrobial Activity. Antibiotics (Basel) 11.

22. Zhou J, Sayre DA, Zheng Y, Szmacinski H, Sintim HO. 2014. Unexpected complex formation between coralyne and cyclic diadenosine monophosphate providing a simple fluorescent turn-on assay to detect this bacterial second messenger. Anal Chem 86:2412–20.

23. Jumper J, Evans R, Pritzel A, Green T, Figurnov M, Ronneberger O, Tunyasuvunakool K, Bates R, Žídek A, Potapenko A, Bridgland A, Meyer C, Kohl SAA, Ballard AJ, Cowie A, Romera-Paredes B, Nikolov S, Jain R, Adler J, Back T, Petersen S, Reiman D, Clancy E, Zielinski M, Steinegger M, Pacholska M, Berghammer T, Bodenstein S, Silver D, Vinyals O, Senior AW, Kavukcuoglu K, Kohli P, Hassabis D. 2021. Highly accurate protein structure prediction with AlphaFold. Nature 596:583–589.

24. Pettersen EF, Goddard TD, Huang CC, Meng EC, Couch GS, Croll TI, Morris JH, Ferrin TE. 2021. UCSF ChimeraX: Structure visualization for researchers, educators, and developers. Protein Sci 30:70–82.

25. Patel DT, O’Bier NS, Schuler EJA, Marconi RT. 2021. The Treponema denticola DgcA protein (TDE ORF 0125) is a functional diguanylate cyclase. Pathogens and Disease doi:10.1093/femspd/ftab004.

26. Vincent AT, Schiettekatte O, Goarant C, Neela VK, Bernet E, Thibeaux R, Ismail N, Mohd Khalid MKN, Amran F, Masuzawa T, Nakao R, Amara Korba A, Bourhy P, Veyrier FJ, Picardeau M. 2019. Revisiting the taxonomy and evolution of pathogenicity of the genus Leptospira through the prism of genomics. PLoS Negl Trop Dis 13:e0007270.

27. Witte G, Hartung S, Büttner K, Hopfner KP. 2008. Structural biochemistry of a bacterial checkpoint protein reveals diadenylate cyclase activity regulated by DNA recombination intermediates. Mol Cell 30:167–78.

28. Bai Y, Yang J, Zhou X, Ding X, Eisele LE, Bai G. 2012. Mycobacterium tuberculosis Rv3586 (DacA) is a diadenylate cyclase that converts ATP or ADP into c-di-AMP. PLoS One 7:e35206.

29. Zheng C, Wang J, Luo Y, Fu Y, Su J, He J. 2013. Highly efficient enzymatic preparation of c-di-AMP using the diadenylate cyclase DisA from Bacillus thuringiensis. Enzyme Microb Technol 52:319–24.

30. Agranoff DD, Krishna S. 1998. Metal ion homeostasis and intracellular parasitism. Mol Microbiol 28:403–12.

31. Toma C, Okura N, Takayama C, Suzuki T. 2011. Characteristic features of intracellular pathogenic Leptospira in infected murine macrophages. Cell Microbiol 13:1783–92.

32. Zhang L, Zhang C, Ojcius DM, Sun D, Zhao J, Lin X, Li L, Li L, Yan J. 2012. The mammalian cell entry (Mce) protein of pathogenic Leptospira species is responsible for RGD motif-dependent infection of cells and animals. Molecular Microbiology 83:1006–1023.

33. Manikandan K, Sabareesh V, Singh N, Saigal K, Mechold U, Sinha KM. 2014. Two-step synthesis and hydrolysis of cyclic di-AMP in Mycobacterium tuberculosis. PLoS One 9:e86096.

34. Putignano V, Rosato A, Banci L, Andreini C. 2018. MetalPDB in 2018: a database of metal sites in biological macromolecular structures. Nucleic Acids Res 46:D459–d464.

35. Tosi T, Hoshiga F, Millership C, Singh R, Eldrid C, Patin D, Mengin-Lecreulx D, Thalassinos K, Freemont P, Gründling A. 2019. Inhibition of the Staphylococcus aureus c-di-AMP cyclase DacA by direct interaction with the phosphoglucosamine mutase GlmM. PLoS Pathog 15:e1007537.

36. Jackson-Litteken CD, Ratliff CT, Kneubehl AR, Siletti C, Pack L, Lan R, Huynh TN, Lopez JE, Blevins JS. 2021. The Diadenylate Cyclase CdaA Is Critical for Borrelia turicatae Virulence and Physiology. Infect Immun 89.

37. Zhukova A, Fernandes LG, Hugon P, Pappas CJ, Sismeiro O, Coppée JY, Becavin C, Malabat C, Eshghi A, Zhang JJ, Yang FX, Picardeau M. 2017. Genome-Wide Transcriptional Start Site Mapping and sRNA Identification in the Pathogen Leptospira interrogans. Front Cell Infect Microbiol 7:10.

38. Pathania M, Tosi T, Millership C, Hoshiga F, Morgan RML, Freemont PS, Gründling A. 2021. Structural basis for the inhibition of the Bacillus subtilis c-di-AMP cyclase CdaA by the phosphoglucomutase GlmM. J Biol Chem 297:101317.

39. Rismondo J, Gibhardt J, Rosenberg J, Kaever V, Halbedel S, Commichau FM. 2016. Phenotypes Associated with the Essential Diadenylate Cyclase CdaA and Its Potential Regulator CdaR in the Human Pathogen Listeria monocytogenes. J Bacteriol 198:416–26.

